# BAMscale: quantification of DNA sequencing peaks and generation of scaled coverage tracks

**DOI:** 10.1101/669275

**Authors:** Lorinc S. Pongor, Jacob M. Gross, Roberto Vera Alvarez, Junko Murai, Sang-Min Jang, Hongliang Zhang, Christophe Redon, Haiqing Fu, Shar-Yin Huang, Bhushan Thakur, Adrian Baris, Leonardo Marino-Ramirez, David Landsman, Mirit I. Aladjem, Yves Pommier

## Abstract

BAMscale is a one-step tool that processes DNA sequencing datasets from chromatin binding (ChIP-seq) and chromatin state changes (ATAC-seq, END-seq) experiments to DNA replication data (OK-seq, NS-seq and replication timing). The outputs include normalized peak scores in text format and scaled coverage tracks (BigWig) which are directly accessible to data visualization programs. BAMscale (available at https://github.com/ncbi/BAMscale) effectively processes large sequencing datasets (~100Gb size) in minutes, outperforming currently available tools.

## Main

Improved technologies and decreasing sequencing costs enable in-depth analysis of chromatin changes for genome-wide comparisons. These studies identify genomic regions with enrichment of binding proteins (ChIP-seq), DNA accessibility (ATAC-seq), DNA breaks (END-seq) or genome replication origin mapping (OK-seq, NS-seq) and timing summarized in Fig.1A-D. In most cases, peak strengths are quantified and normalized by performing multiple analysis steps. This is usually carried out with “in-house” scripts i.e. time-consuming case-by-case programming. Although there are available tools for sequencing track generation [1–3], they either require multiple steps, and/or need more computation time to generate results ready for visualization.

**Figure 1.**
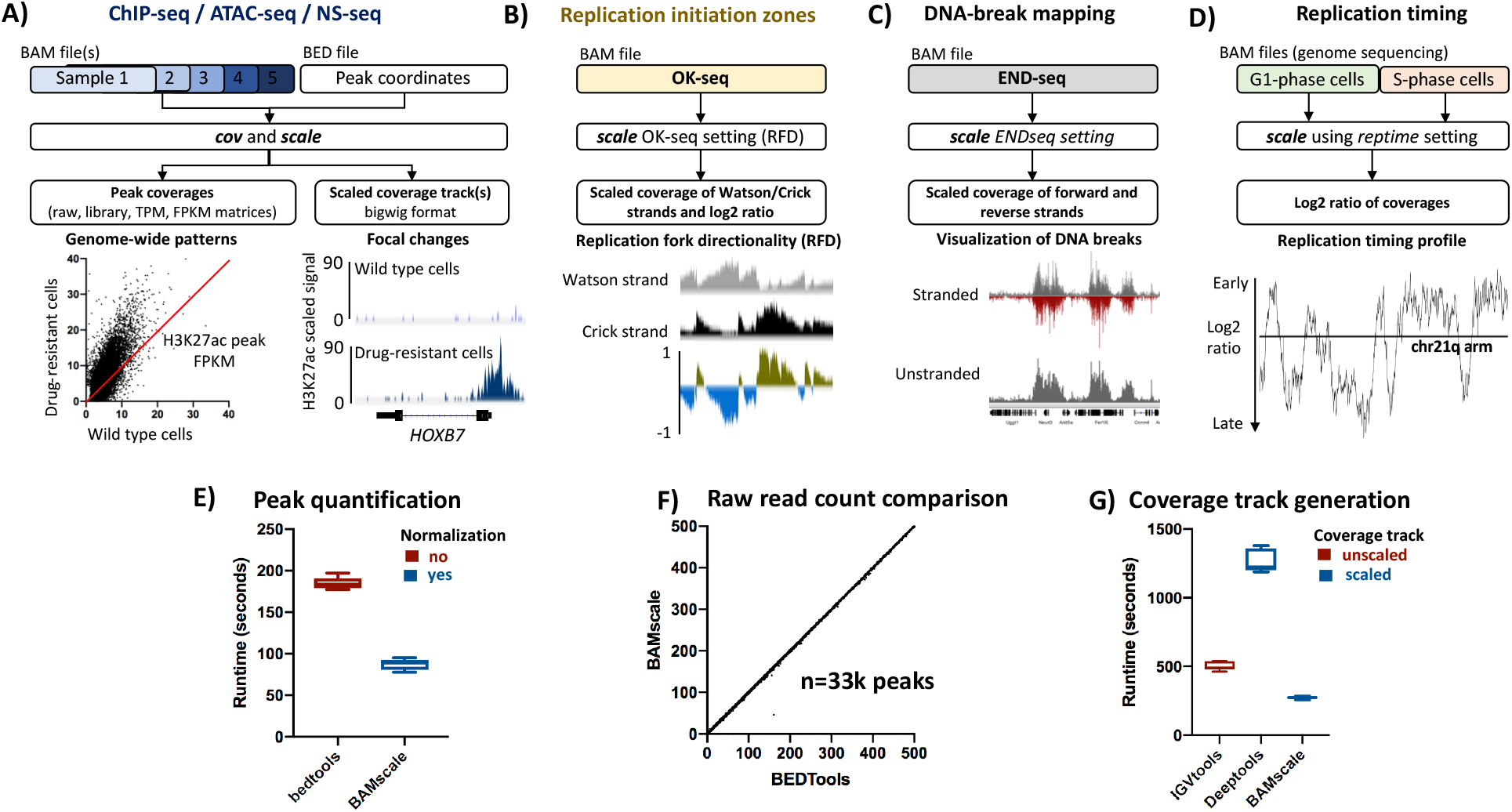
Application and benchmarking of BAMscale on different sequencing datasets. A) Scaled coverage track generation and peak quantification of ChIP-seq and ATAC-seq data. Local differential H3K27ac signal at the HOXB7 locus in MV4-11 (wildtype) and PKC412-resistant (R) (drug resistant) cells, and global H3K27ac increase. B) OK-seq coverage tracks can be generated in one step, outputting scaled strand-specific coverage tracks, and the replication-fork directionality (red). C) Mapping DNA-breaks from END-seq, creating strand-specific or unstranded coverage tracks. D) Analysis of replication timing data. E) Performance comparison of peak quantification and F) correlation of raw read counts in ~33k peaks between *BAMscale* and *bedtools*. G) Coverage tracks generation benchmarks using IGVTools (unscaled output), *Deeptools* and *BAMscale* (scaled output).

Here we introduce BAMscale, a new genomic software tool for generating normalized peak coverages and scaled sequencing coverage tracks in BigWig format. These two functions enable rapid and highly accurate identification of genome-wide and local changes by normalizing and scaling data in a single step. To achieve higher accuracy in peak quantification, our tool by default calculates the number of aligned reads by processing the entire BAM file, producing better alignment estimates than by simply using the aligned BAM index file where duplication metrics are not present. This feature is important in cases where a set of samples have higher duplication rates, skewing the results of normalization (**supplementary methods**). The default coverage track generation involves normalizing the per-base (binned) coverage to the total number of aligned bases divided by the genome size. Since our tool is developed in the C language using the *samtools* library [4], it has superior performance to existing software. BAMscale can process ~100GB of aligned data (in BAM format) in under 20 minutes using a computer with 4 processing threads.

BAMscale quantifies ChIP-seq/ATAC-seq peaks from BAM and BED files producing raw read counts, as well as TPM, FPKM and library size normalized peak scores (Fig.1A). By providing accurate peak quantification in parallel with generated scaled coverage tracks, BAMscale simplifies the comparison and visualization of genome-wide and local changes. To illustrate this point, we reanalyzed published histone ChIP-seq data from MV4-11 cell line and their PKC412 (a multi-target protein kinase inhibitor) resistant counterpart (MV4-11R) [5]. In agreement with published results, we observed a global increase of H3K27ac, a decrease in H3K27me3 and a predominantly unchanged H3K4me3 signal in the drug-resistant cells (Fig.1A, Fig.S1). Drug resistant cells displayed elevated protein expression of HOXB7 [5], which has increased histone H3K27ac signal, a known marker for active genes.

BAMscale also enables processing of Okazaki fragment sequencing (OK-seq) data in a single step, generating scaled strand-specific coverage tracks (of Watson and Crick strands), as well as quantified replication fork directionality (RFD) ratios, as shown for K562 cells [6] (Fig.1B). OK-seq identifies genomic regions where replication origins have synchronized initiation and directionality [7]. In this approach, RNA-primed Okazaki fragments are measured using strand-specific sequencing. Genomic regions with synchronized origins display a shift in positive and negative (Watson/Crick) strand ratios.

Replication-timing sequencing involves identifying copy-number state differences between G1-phase and replicating S-phase (or asynchronous - AS) cells. The G1-synchronized cells have a diploid genome state (2N copies) while copy-number status of replicating cells ranges between 2N for late-replicating regions and 4N for early-replicating regions [8, 9]. The results of replication-timing sequencing consist of two (or more) genome sequencing files with high coverages (usually >50x), which are used to classify and identify the replication timing of the genome. BAMscale processes and generates scaled-coverage tracks, as well as the log_2_ coverage ratios for the entire human genome in minutes (Fig.1C). Additionally, BAMscale includes a simple script to generate BED formatted segments of early-, mid-early-, mid-late- and late-replicating regions.

While performance of BAMscale for peak quantification is comparable to the most commonly used *bedtools[2]* program using a single processing thread, BAMscale reduces execution time to ~50% when using multiple threads (Fig.1D). In addition, *bedtools* will only calculate raw read counts, while BAMscale performs normalization of raw read counts while outputting FPKM, TPM and library size normalized peak scores. This enables a direct comparison of peaks between conditions. Correlation of raw read counts from the two methods is above 0.99 (Fig.1E and Fig.S2).

Most studies employing DNA-capture-based sequencing methods concentrate on local examples with genes of interest. Current popular methods either generate unscaled coverage tracks or require multiple processing steps and computational time. BAMscale is capable of generating scaled coverage tracks, enabling a direct comparison of signal intensities using the IGV[3] browser or the UCSC genome browser [10]. BAMscale outperforms the popularly used IGVtools using one (or multiple threads) by over 1.5-fold (Fig.1F) in track generation. Additionally, IGVtools computes unscaled coverage tracks with no possibility for read filtering (such as duplicate reads, or poor alignment quality). The execution time of BAMscale is approximately 6-times quicker than the *deeptools*[1] *bamCompare* program (Fig.1E) for scaling coverage tracks. This is important when large BAM files have to be processed, such as replication timing data, where BAMscale can create scaled coverage tracks and log_2_ coverage ratios for ~100Gb of data in approximately 20 minutes. BAMscale can be easily used to compare multiple different datasets, such as OK-seq, END-seq and replication timing, generating reproducible results [11] (Fig.S3-S4).

To further demonstrate the potential of BAMscale, we compared differences in chromatin accessibility from ATAC-seq data recently published by our group [12]. In the analysis, we compared the effect of camptothecin (CPT) treatment in CEM-CCRF (*SLFN11*-positive) cells and their isogenic *SFLN11*-knockout. After CPT treatment, chromatin accessibility remained unchanged in the *SLFN11*-KO cells, while accessibility of pre-existing sites strongly increased in the *SLFN11*-positive cells (Fig.2A). Using the GIGGLE tool [13] on the Cistrome [14] website, we found that ATAC-seq peaks strongly overlapped H3K27ac, H3K4me3 and H3K9ac sites, which are histone marks associated with active genes (Fig.2B). Colocalization analysis of sites with >3-fold increase during CPT treatment in *SLFN11*-positive cells showed ~20% increase in overlap with H3K4me3 and H3K9ac sites, identified using Coloweb [15] (Fig.S5, supp. table 1). DNA accessibility sites were strongly enriched in gene promoter regions, such as in the *TOP1* and *CTCF* gene promoters (Fig.2C).

**Figure 2.**
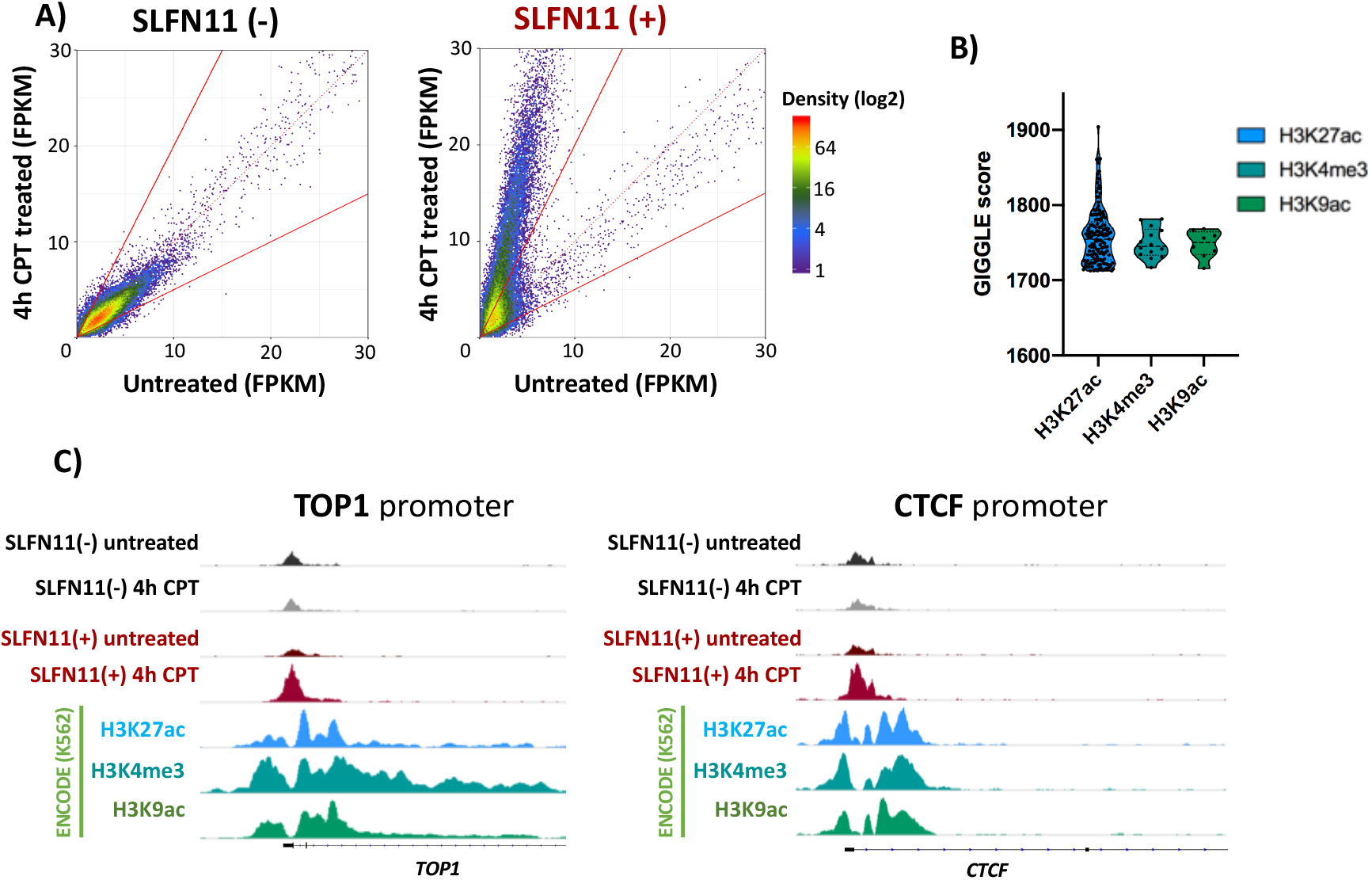
Application of BAMscale on ATAC-seq data. A) ATAC-seq signal change is observed in wild-type CEM-CCRF cells (*SLFN11* positive), and not in the *SLFN11* isogenic knockout. B) Colocalization of opening ATAC-seq peaks using GIGGLE and cistrome. C) Examples of chromatin accessibility in the *TOP1* and *CTCF* genes.

Next, we compared replication timing data to OK-seq and NS-seq (Nascent strand sequencing) in the human leukemia K562 cell line. Replication timing results (Fig.3A i) and the generated segments (Fig.3A ii) showed that early-replicating regions strongly correlate with active chromatin (Fig.3A iii) identified with ChromHMM [16, 17]. Furthermore, BAMscale showed a strong overlap of OK-seq [6] RFD strand switches (associated with synchronized replication initiation zones) with active (eu)chromatin (Fig.3A iv,v). Fewer than 0.5% of identified OK-seq strand switches were identified in heterochromatin, where no overlap with active chromatin regions was found. Similarly, we observed higher NS-seq signal (and replication origin peaks) in euchromatin (Fig.3A vi). Early-replicating regions tend to have more replication initiation sites, which gradually decrease in later phases of replication timing (Fig.3B). These results correlate strongly with the NS-seq results showing that early replicating regions have higher peak densities compared to later-replicating regions (Fig.3C).

**Figure 3.**
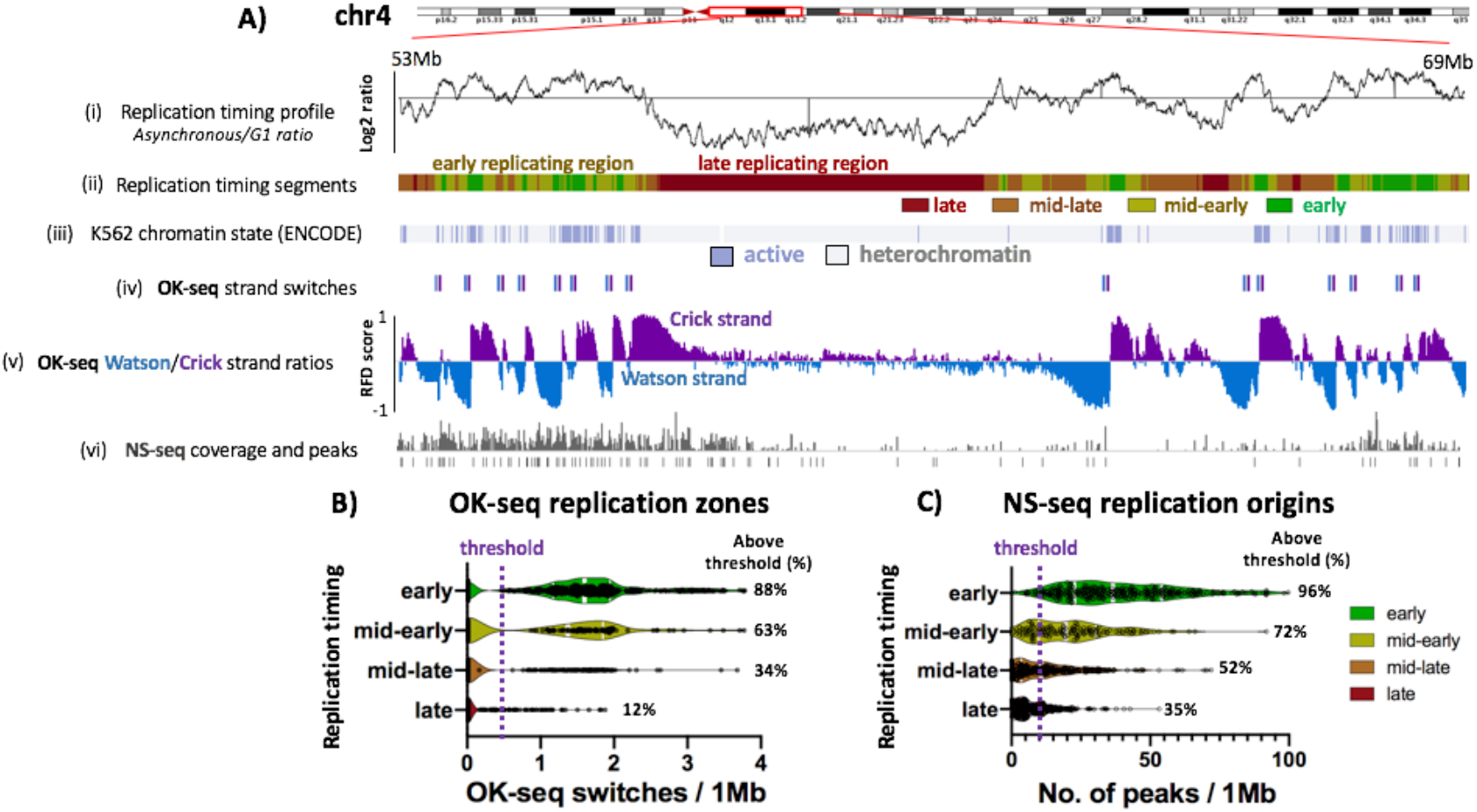
Comparison of different replication sequencing methods. A) Replication timing ratios (i), replication timing profile (ii), active/repressed chromatin regions (iii), strong OK-seq strand switch coordinates (iv), OK-seq replication-fork directionality ratios (v) and NS-seq (replication origin) tracks for K562 cell line. B) OK-seq strand switches in the four segments of replication. C) NS-seq peak abundances in the four replication timing phases.

Widespread usage of DNA capture-based methods helps us understand and categorize changes in chromatin state and their regulatory effects. Using BAMscale as a peak quantification method and a scaled coverage-track generation tool, we are capable of identifying single focal changes on the genome as well as understanding how certain conditions alter the chromatin profile. Finally, to our knowledge, BAMscale is the only tool that can directly output scaled stranded (Watson/Crick) coverages and RFD tracks for visualization of OK-seq data and stranded coverage tracks for END-seq data.

## Acknowledgements

This research was supported by the Intramural Research Programs at the National Cancer Institute and the National Library of Medicine at the NIH. The study utilized the high-performance computer capabilities of the Biowulf HPC cluster at the NIH.

## Data availability

Raw sequencing data and reprocessed files are available from NCBI under accession GSE131417.

## SUPPLEMENTARY FIGURES

**Supplementary figure 1.**
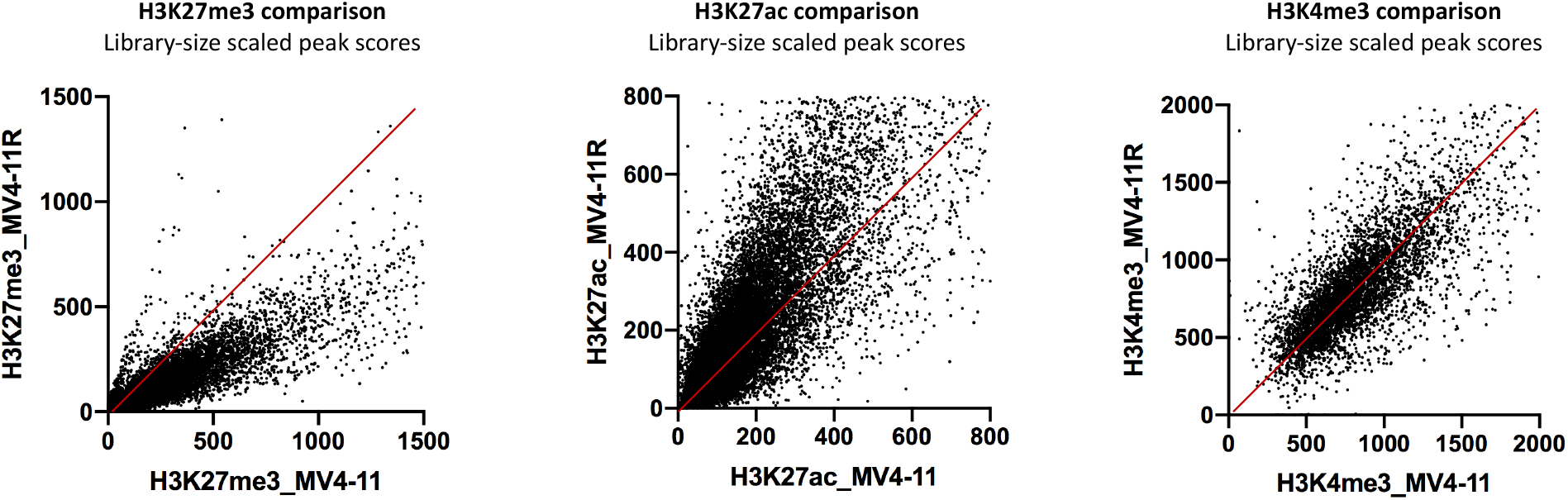
Comparison of three histone marks in MV4-11 and MV4-11R cells.

**Supplementary figure 2.**
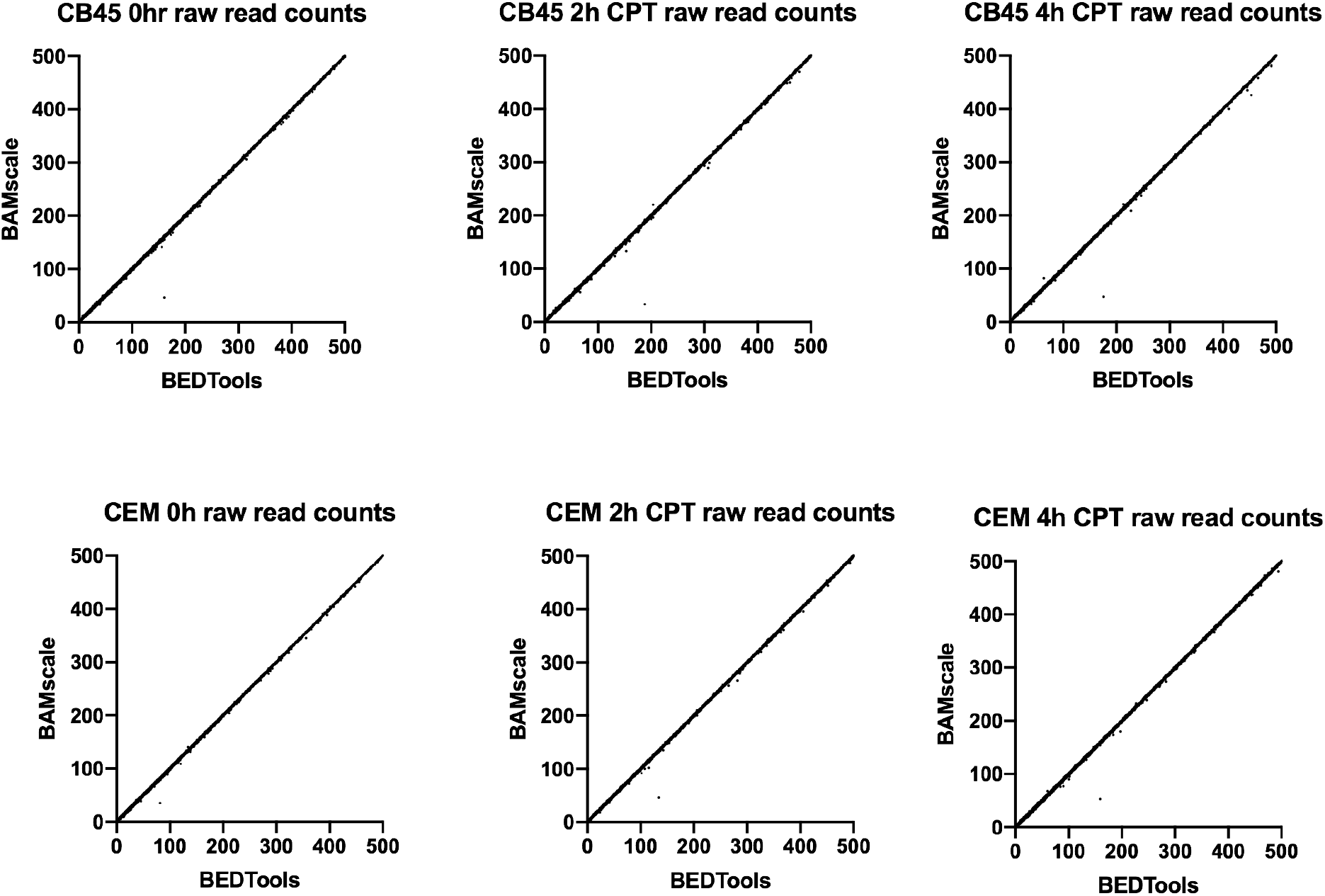
Comparison of raw read counts analyzed with BAMscale and BEDtools.

**Supplementary figure 3.**
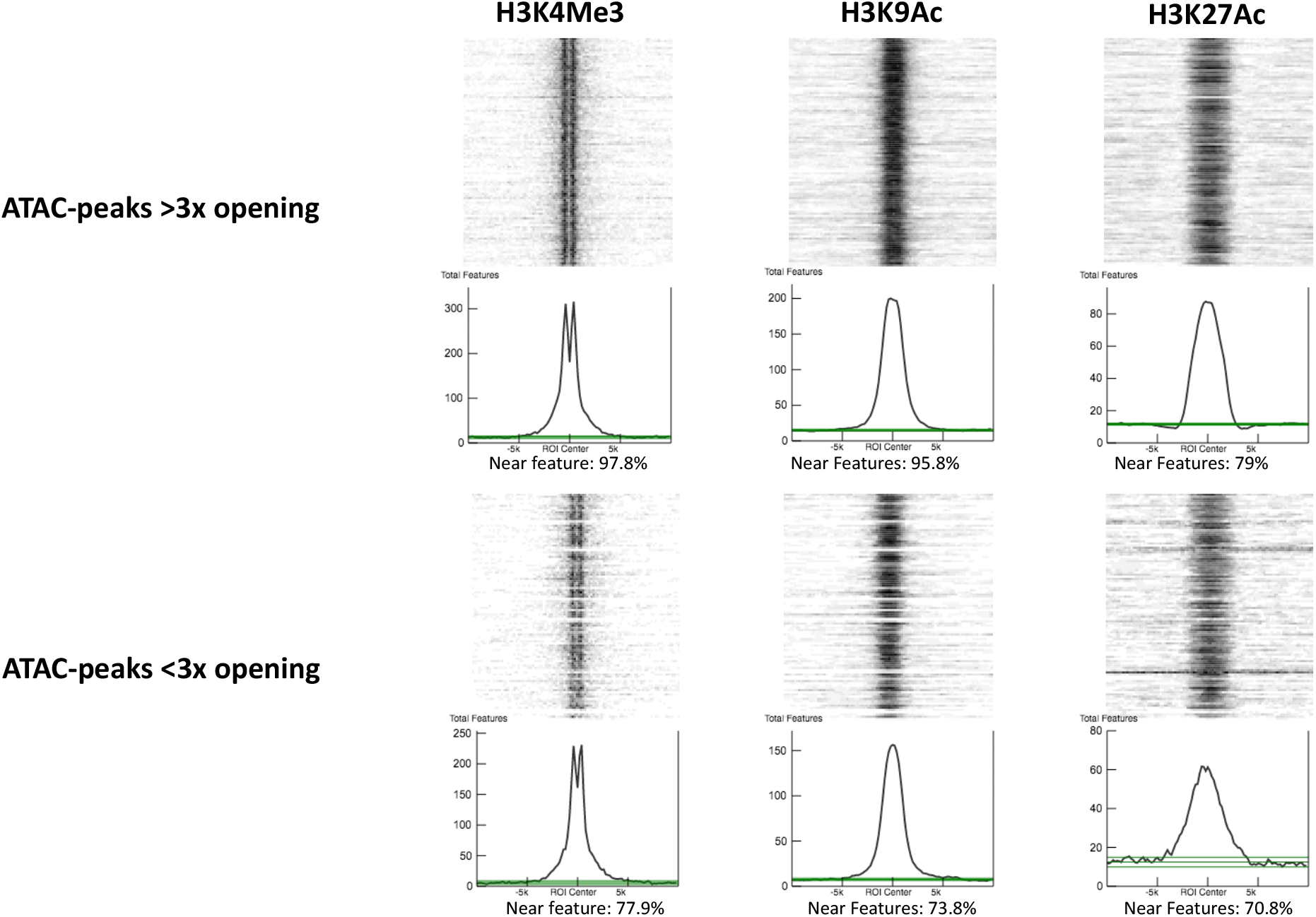
Colocalization of ATAC-seq peaks and H3K4me3, H3K9ac and H3K27ac marks.

**Supplementary figure 4.**
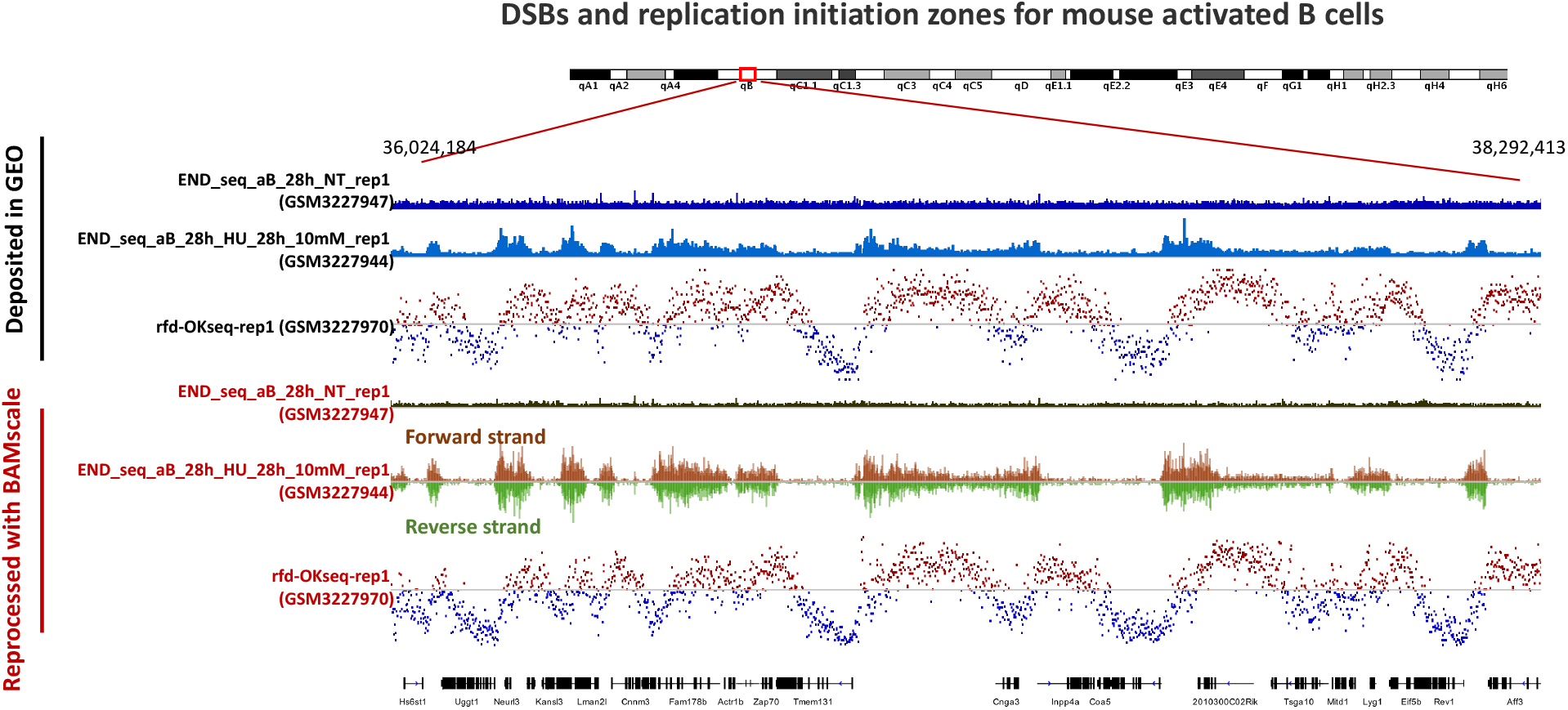
Comparison of deposited and reprocessed END-seq and OK-seq tracks.

**Supplementary figure 5.**
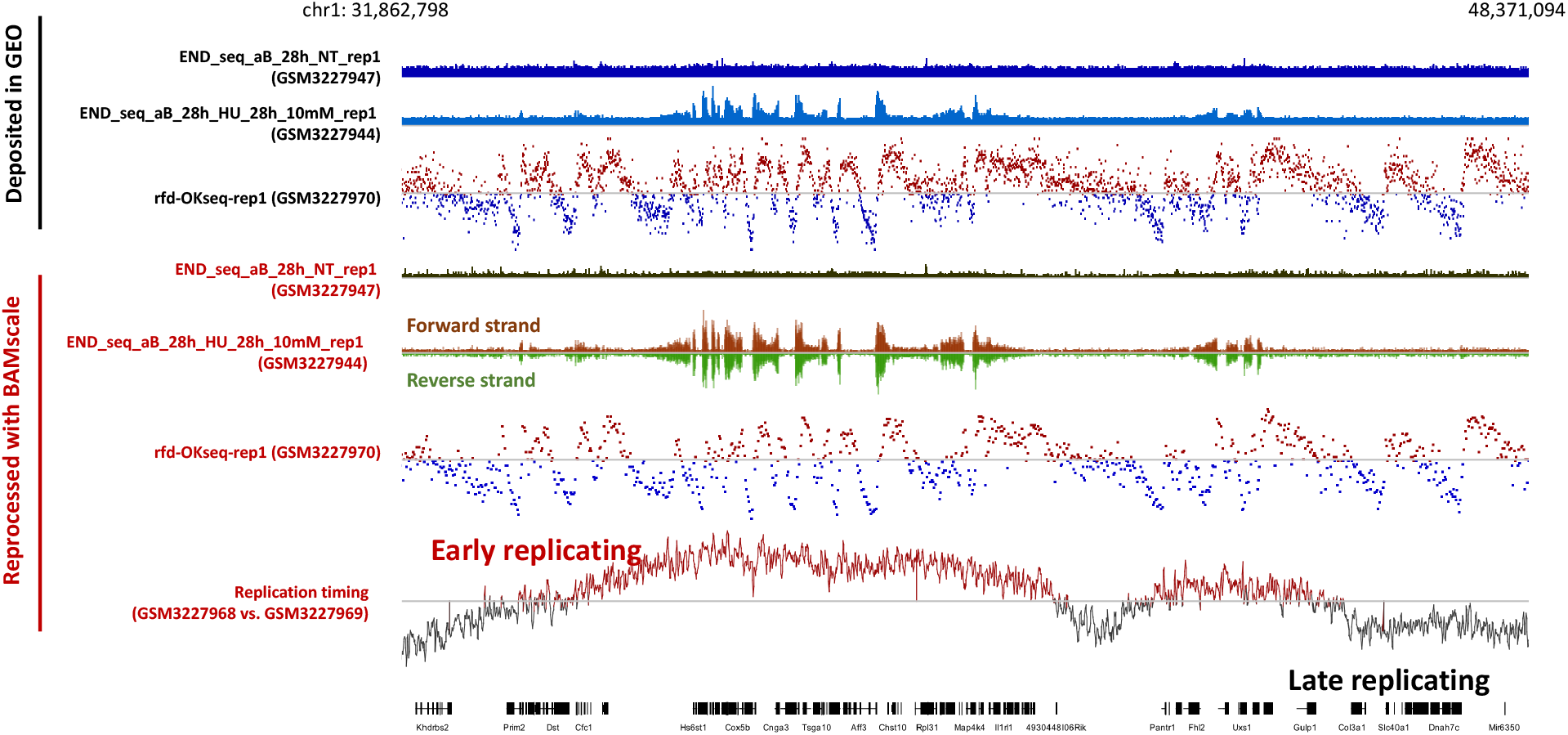
Comparison of deposited and reprocessed END-seq and OK-seq tracks showing replication timing.

## SUPPLEMENTARY TABLES

**Supplementary table 1.** Colocalization statistics of ATAC peaks with histone marks from U2OS cells calculated with Coloweb.

| Colocalization of peaks >3x opening from CPT treatment (Coloweb, K562, ROI centered) |  |  |  |  |  |  |
| --- | --- | --- | --- | --- | --- | --- |
| Feature | AMI | BMI | Peak Height | Percentage Near Features | Total Features | Average Feature Length |
| H3K4Me3 | 2974.8 | 0 | 303.4 | 97.80% | 58775 | 151 |
| H3K9Ac | 2486.4 | 0.5 | 185 | 95.80% | 36409 | 991 |
| H3K27Ac | 1186.6 | 13.5 | 76 | 79.00% | 58937 | 2783 |
| H3K79Me2 | 1174.6 | 2 | 63 | 40.00% | 29111 | 9625 |
| H3K4Me2 | 903.7 | 23.6 | 52 | 74.10% | 70379 | 2086 |
| H3K4Me1 | 729.8 | 0.2 | 48.9 | 95.70% | 108851 | 627 |
| H4K20Me1 | 650 | 3.7 | 34.3 | 42.30% | 44202 | 18011 |
| H3K9Me1 | 106.1 | 8.8 | 11.4 | 61.60% | 88350 | 8569 |
| H3K9Me3 | 54.7 | 0.2 | 9.4 | 25.20% | 43231 | 21534 |
| H3K36Me3 | 45.2 | 245.2 | 11 | 53.30% | 36411 | 151 |
| H3K27Me3 | 12.3 | 0 | 17.4 | 0.90% | 1379 | 151 |
| Colocalization of peaks <3x opening from CPT treatment (Coloweb, K562, ROI centered) |  |  |  |  |  |  |
| Feature | AMI | BMI | Peak Height | Percentage Near Features | Total Features | Average Feature Length |
| H3K4Me3 | 2201.2 | 0 | 224.9 | 77.90% | 58775 | 151 |
| H3K9Ac | 1894.6 | 0.3 | 148.7 | 73.80% | 36409 | 991 |
| H3K27Ac | 1000.5 | 0 | 67.7 | 70.80% | 58937 | 2783 |
| H3K79Me2 | 851.7 | 0 | 49.2 | 35.50% | 29111 | 9625 |
| H3K4Me2 | 777.5 | 0.6 | 47.3 | 68.20% | 70379 | 2086 |
| H3K27Me3 | 756.7 | 0 | 214 | 2.90% | 1379 | 151 |
| H3K4Me1 | 661.7 | 2.5 | 43.5 | 77.00% | 108851 | 627 |
| H4K20Me1 | 660.4 | 4.2 | 31.8 | 39.50% | 44202 | 18011 |
| H3K9Me1 | 162.4 | 0.1 | 10.5 | 57.10% | 88350 | 8569 |
| H3K9Me3 | 98.9 | 0 | 10.7 | 29.10% | 43231 | 21534 |
| H3K36Me3 | 3.4 | 24.2 | 7.1 | 37.80% | 36411 | 151 |

